# Photo-Uncaging Triggers on Self-Blinking to Control Single-Molecule Fluorescence Kinetics for Super-Resolution Imaging

**DOI:** 10.1101/2024.02.13.580074

**Authors:** Ying Zheng, Zhiwei Ye, Xue Zhang, Yi Xiao

**Affiliations:** State Key Laboratory of Fine Chemicals, Frontiers Science Center for Smart Materials Oriented Chemical Engineering, Dalian University of Technology, Dalian 116024, China; Dalian Medical University, Dalian 116000, China

**Keywords:** self-blinking rhodamines, super-resolution image

## Abstract

Super-resolution imaging in a single-molecule localization approach has transformed the bulk fluorescence requirements to a single-molecule level, raising a revolution in the fluorophore engineering. Yet, it is a challenge to structurally devise fluorophores manipulating the single-molecule blinking kinetics. In this pursuit, we have developed a new strategy by innovatively integrating the photoactivatable nitroso-caging strategy into self-blinking sulfonamide, to forming a nitroso-caged sulfonamide rhodamine (NOSR). Our fluorophore demonstrated controllable self-blinking events upon photo-triggered uncaging release. This exceptional blink kinetics improved integrity in super-resolution imaging microtubules compared to self-blinking analogues. With the aid of paramount single-molecule fluorescence kinetics, we successfully reconstructed the axial morphology of mitochondrial outer membranes. We foresee that our synthetic approach of photoactivation and self-blinking would set a new avenue for devising rhodamines for super-resolution imaging.

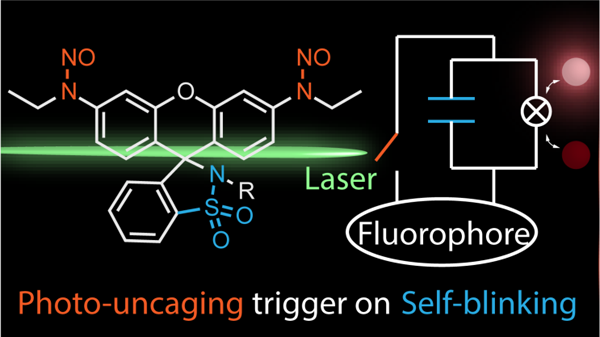

## Introduction

Super-resolution imaging on the basis of single-molecule localization has unambiguously pushed the boundaries of life science to a molecular level.^1,2^ The resolution enhancement capitalizes on the precise location recovery of each fluorophore molecules through blinking events, i.e. spatial and temporal distributed dark-to-bright events in a sparse fashion.^3-5^ This molecular localization mechanism pushes the fluorophore requirement to a single-molecule level beyond conventional ensemble conditions. Although molecular structure fundamentally impacts the blinking kinetics, the engineering of desired fluorophores remains exception-ally rare.^6,7^ Existing strategies confront barriers in obtaining sparse single-molecule fluorescence in high-content labeling of structure determinations (e.g., immunostaining labeling). There is a far way before reaching limits in devising fluorophores with controllable blinking kinetics for localization based super-resolution imaging.

Recent years have witnessed significant progression in modifying single-molecule blinking kinetics of rhodamines. Successful cases including photoactivation through caging/photoreaction strategy,^8-14^ self-blinking introduced by the spirocyclization equilibrium,^15-19^ or stabilizing strategy through hydrogen bonds^20-22^. The photoreaction strategy completely conceals the fluorophores into non-emissive states through light-controlling mechanics. Prominent cases including 2-diazoketone caged rhodamine,^23,24^ PaX dyes,^25^ and nitroso-caged rhodamine.^26,27^ Self-blinking strategy develops an exquisite spirocyclization equilibrium through designed substitution on the 9 position of xanthene. This strategy was first elucidated by Urano group by a landmark Si-rhodamine derivative (HMSiR),^28,29^ and later explored by different groups. ^30-35^ For the stabilizing strategy, hydrogen-bonding functional groups were directly introduced to the fluorophore scaffold, improving the stability of zwitterionic open-ring or leuco close-ring structure.^20-22^

However, these strategies fail to tune the entire spectrum of the single-molecule blinking kinetics. The self-blinking events are the intrinsic kinetics of ring-opening process, depended on the fluorophore structure. This structural nature narrows the application window of self-blinking dyes, as their blink kinetics show limited flexibility. The photo-reaction strategy adds light control utility for single-molecule bright events, yet it does not modulate the blink kinetics of the photoproducts, resulting in none sufficient localizations. The stabilizing strategy adjusts the kinetics of dark-bright transform as well as the ratio of bright molecules. But the strategy provides no control for the dark-bright blink events. Thus, obtaining ideal single-molecule blinking kinetics through molecular structure design remains a huge challenge.

In general, self-blinking strategy and caging modifications, constitute two complementary approaches for con-trolling the blinking kinetics, as the former provides intrinsic dark-bright events and the latter enables photo-triggered control. The advantages of self-blinking strategy and caging strategy cover each other’s disadvantages. If properly integrated, the caging modification grants the controlling of self-blinking events in a photo-gated fashion. Thus, introducing the caging strategy into the self-blinking strategy would controllably tune the single-molecule blinking kinetics from a structural perspective. However, it is a challenge to combine these strategies and no attempts have been per-formed priorly. Traditional o-nitrobenzyl caged rhodamine requires complex synthesis and bio-toxic UV activation light.36 2-diazoketone caging37,38 as well as the caging-group-free strategy25,39,40 required modifications or removes of rhodamine spiro ring, raising issues for developing self-blinking thermal equilibrium on the basis of spirocyclization. There is a great barrier for introducing caging strategy into the self-blinking strategy in controlling single-molecule blinking kinetics.

Recently, we have developed a visible-light photoactivatable nitroso-caged strategy, with a small nitroso unit deco-rating the donor amine groups.^26^ This minimalist unit does not interfering the cyclization equilibrium of the rhodamine spirolactams, facilely introducible to the self-blinking fluorophores. Thus, we designed and synthesized nitroso-caged sulfonamide rhodamine (NOSR, Figure 1a) through the craft of a new photo-trigger self-blinking strategy. In single-molecule studies, NOSR exhibited a large number of blinking events and controllable single-molecule dark-bright transform. The photo-triggered strategy surpassed the shortage of narrow imaging time window of sulfonamide rhodamine, without disturbing its rapid self-blinking kinetics (krc = 0.6-4.6 s^-1^), expanding the imaging adaptivity to high-density labeling immunostaining condition. Finally, the super-resolution imaging capability of NOSR was studied, demonstrating the efficacious of molecular engineering for blink kinetics.

**Figure 1.**
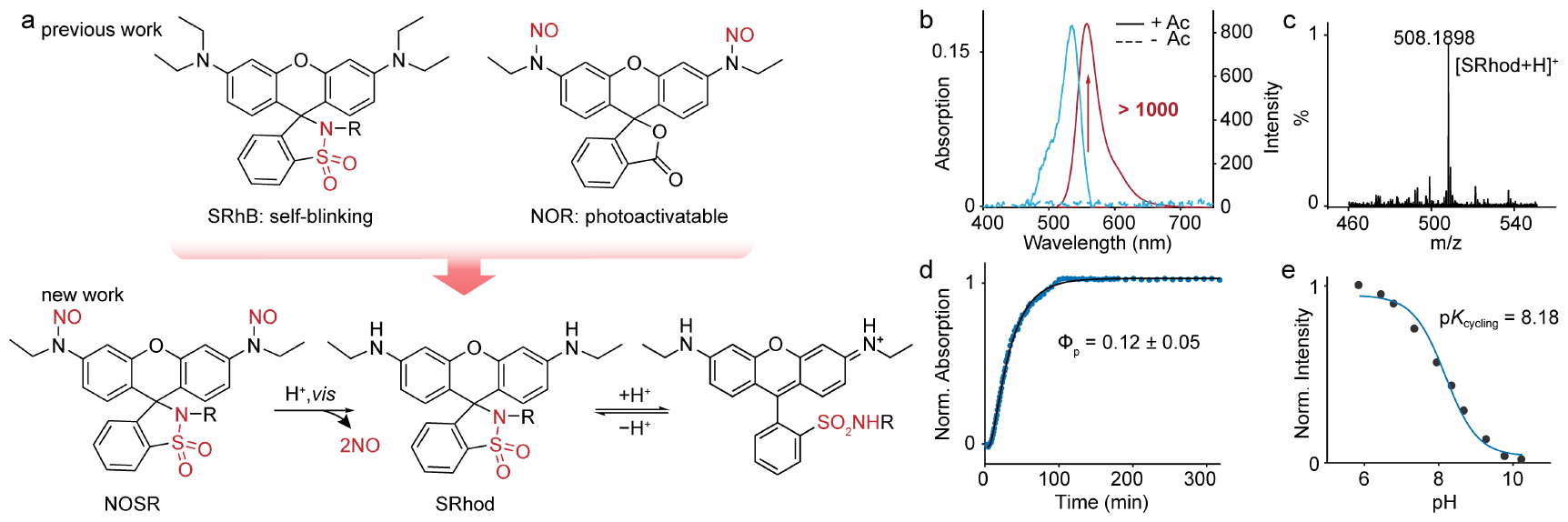
Chemistry of photo-triggering upon self-blinking strategy. (a) The molecular design strategy of NOSR. (b) Spectra of NOSR before and after photoactivation. (c) High-resolution mass spectrometry of the photoproducts of NOSR. (d) exhibit the acquired absorption data at peak wavelength (535 nm for NOSR, blue dots). (e) Comparison on pH titration curves of the photoproducts of NOSR.

We first investigated the spectral properties of NOSR. In vitro spectral measurement revealed that NOSR shown neither visible absorption nor fluorescence bands, indicating that the caging units successfully forced NOSR to lueco ring-closed form. Upon irradiation with UV/visible light, the absorption (fluorescence) peaked at 535 (558) nm gradually appeared and increased until photoactivation saturation was reached (Figure S1). The generation of visible fluorescence from none demonstrated a high dark-bright ratio (> 1000, Figure 1b) and the resulting photoproducts, sulfonamide rhodamine (evidenced by a high-resolution mass spectrum m/z at 508.1898, Figure 1c), exhibited high brightness (Φ = 0.83, ε = 96000 L·mol^-1^·cm^-1^). NOSR displayed a primary photoconversion quantum yield (ca. 0.12 ± 0.05) comparable to classic nitroso-caged rhodamine^26^ (Figure 1d). More importantly, the photoproducts SRhod exhibited a spirocyclization equilibrium with p*K*_cycling_ = 8.18 (Figure 1e and S2), suggesting a natural dark-bright stochastic switch. NOSR was photoactivated to a photoproduct of high brightness at zwitterionic ring-opening form, capable to switch its fluorescent state through a thermal equilibrium, highlighting its promising potential for localization imaging.

Encouraged by the spectral properties of NOSR, we next studied the single-molecule photophysics. Two self-blinking analogues, SRhod (obtained through complete photoactivation of NOSR) and SRhB, were compared in the study to understand the blink kinetics of NOSR. Figure 2a shows the typical fluorescence trajectories obtained. NOSR exhibits unique photo-gated self-blinking events sparsely controlled throughout the entire imaging process compared to the other two fluorophores. NOSR continually photoactivated in time, and its photoproducts produce dozens of blink events followingly, whereas the majority of SRhod and SRhB molecules provide their blink events at the first ten seconds due to fast intrinsic ring-opening rates. The nitroso-caging strategy shifts the temporal distribution of self-blinking events.

**Figure 2.**
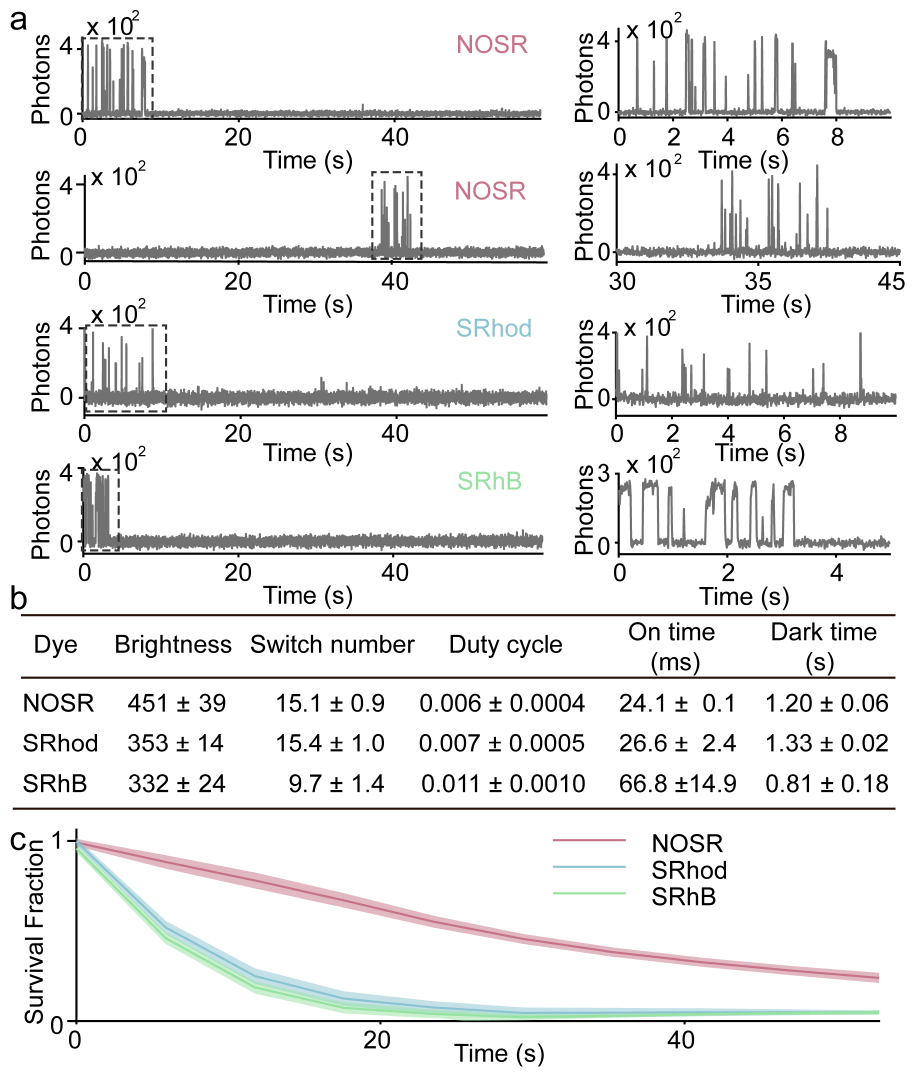
Single-molecule photophysical studies of NOSR, SRhod, and SRhB. Typical fluorescence trajectories (a), the summary of single-molecule characteristics (b) and the molecular survival as a function of time (c) under 532 nm laser (2.0 kW/cm^2^) irradiation.

Figure 2b (Table S1) summarizes the single-molecule photophysics. Since NOSR is photoactivated to SRhod, the two exhibit similar single-molecule switch numbers (∼15), duty cycles (∼0.006-0.007), on times (∼24-27 ms) and dark times (∼1.2-1.3 s). SRhB exhibits slightly prolonged on time (67 ms), reduced switching number (9.7) and shortened dark time (0.8 s) among the three, which might correspond to its extra ethyl substitution. According to the quantitation, trigger-strategy NOSR demonstrates similar single-molecule performance as the other analogues. Then, what is the uniqueness of the blink kinetics of NOSR?

The self-blinking events is redistributed in time for the trigger-strategy fluorophore. The addition of nitroso-caging unit adds a photo-control mechanism on the dark state, regulating the single-molecule signal continuity and sparsity. NOSR demonstrates an over four-fold slower decaying rate in the ratio of survival molecules compared to that of SRhod and SRhB (Figure 2c, *τ*_1/2_: NOSR, 30 s *vs* SRhod, 7 s *vs* SRhB, 6 s). The caging construction delays the formation of bright events, prolonging the temporal window of successive fluorescence signals (whereas the photon-emissive events of other fluorophores might lose during depletion stage, Figure S3). In short, through integration of caging upon self-blinking, NOSR molecule demonstrates photo-gated single-molecule dark-bright events in a temporal sparsity. The prolongation of survival fraction makes NOSR a favorable candidate for high-quality single-molecule localization super-resolution imaging in multi-labeling immunostaining condition.

Next, we investigate the imaging capability of NOSR through immunostaining microtubules and mitochondrial outer membranes. In echo to the controllable blinking kinetics, pure PBS buffer was used in the imaging, excluding any imaging enhancers. NOSR is successfully labelled to target structures (Figure S4-S5), and continuously provides sparse fluorescence signals during imaging. The twisting and bending of microtubule fibers were successfully recon-structed through the localization of NOSR molecules (Figure 3a). Magnified images reveal the significant disparity on clarity between conventional and super-resolution imaging (Figure 3b-c). The super-resolved reconstruction precisely measures the distance of the two microtubule fibers at of 127 nm (Figure 3d). Consistent with microtubule imaging results, NOSR fully reconstructs the morphology of mitochondria (Figure 3e-g). Moreover, reconstruction quantitatively resolves the mitochondria width at 240-249 nm (Figure 3h). The corresponding Fourier ring correlation (FRC)^41,42^ analysis shows a relatively high spatial resolution (microtubule, 93.4 nm; mitochondria, 99.1 nm, Figure S6).

**Figure 3.**
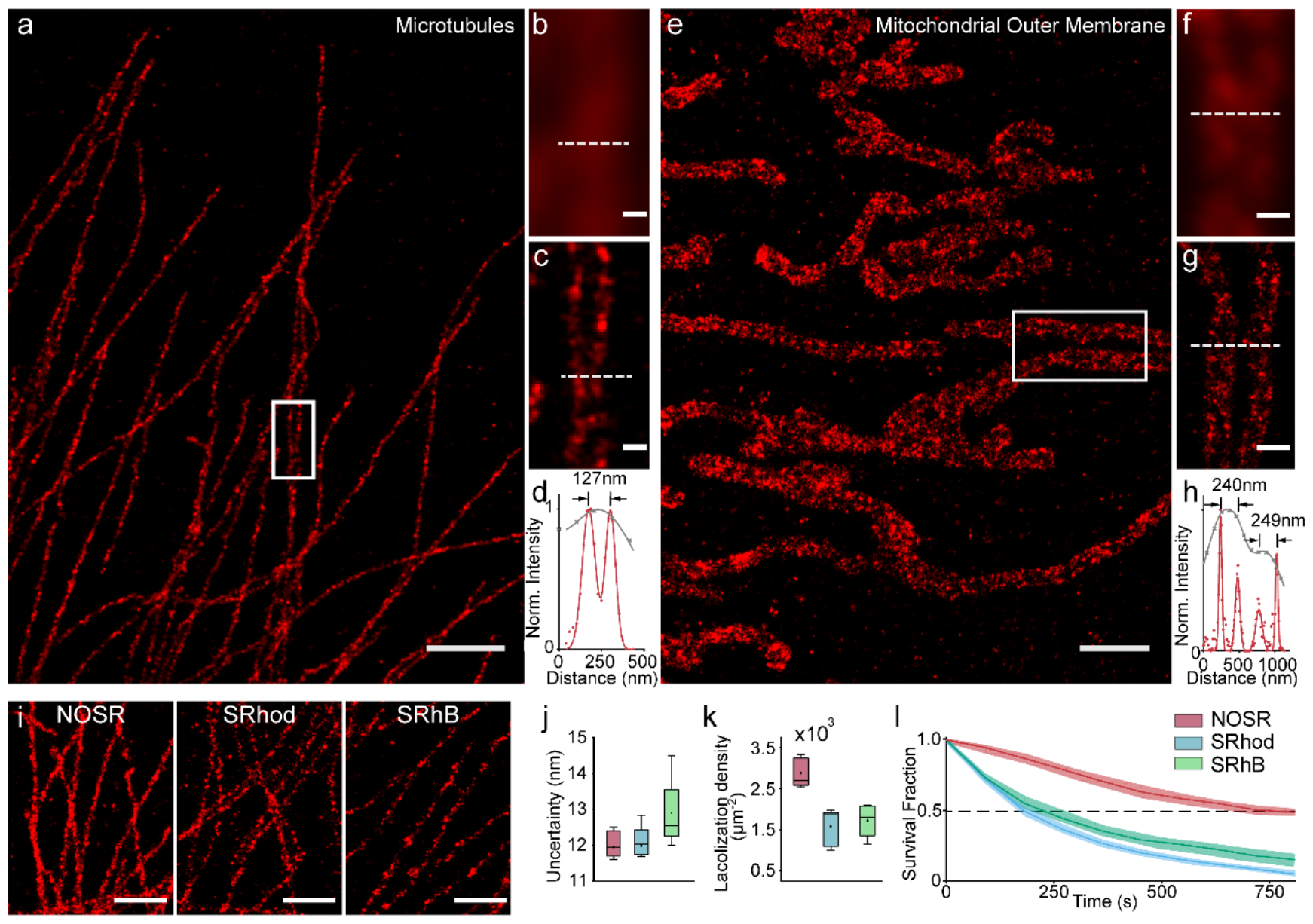
Super-resolution image of microtubules (a) and mitochondrial outer membrane (e) immunolabeled with NOSR. The conventional image (b, f) and super-resolution image (c, g) of a magnified view at the highlighted regions in (a, e). (d, h) Intensity pro-files indicated with white lines in (c, g). (i) Comparison of on the super-resolution imaging of microtubules. The box plots of localization uncertainties (j) and densities (k) on reconstructed microtubules (cell number > 5). (l) Comparison of the molecular survival fraction as a function of time. Scale bars: 2 μm (a); 0.2 μm (b, c); 1 μm (e, i); 0.5 μm (f, g).

To confirm control blinking kinetics in impacting imaging capability, we further compared the super-resolution imaging results between NOSR, SRhod and SRhB under identical conditions (dye-antibody ratio: ∼1.6, Figure 3i and S7). Among the three, NOSR presents an enhanced localization precision (11.8 nm, NOSR vs 12.0-12.8 nm, SRhod, SRhB), and the highest localization density on microtubule fibers (> 2400 localizations·μm^-2^, Figure 3j-3k and S8). The high localization density indicates an enhanced integrity on the microtubule reconstruction, suggesting an imaging supremacy of NOSR. The enhancement is a consequence of temporal shift of the self-blinking events upon photo-gated fashion, as the trigger-strategy NOSR shows blinking kinetics free from the concentrated overlapping signals. Temporal evolvement analysis of re-construction reveals that both SRhod and SRhB sharply arrives signal attenuation under laser irradiation, whereas the signals of NOSR attenuated mildly forming continual structural features (Figure S9-S10). The redistribution of self-blinking events of NOSR was further validated in molecular survival analysis (Figure 3l). NOSR molecules exhibit a survival half time at 695 s, which is 3.0-3.8 times slower than SRhod and SRhB. In summary, the introduction of nitroso-caging groups grafts photoactivation to the self-blinking events, controlling the blinking kinetics to make NOSR efficacious for super-resolution imaging under high-content immunostaining conditions.

The extension of imaging dimensions from two to three further pushes the blink kinetics requirement to an unprecedent level. In three-dimensional imaging, morphology of the point spread function (single-molecule signal) is utilized for extracting axial location, and any background or overlapping between signals would completely distort the results.^43-45^ The trigger-strategy fluorophore controllably generates single-molecule signals sufficiently sparse for axial localization. Through axial position recovery, the three-dimensional imaging of mitochondrial outer membrane and microtubules is successfully reconstructed (Figure 4 and S11). The x-z projection view of mitochondria shows a complete circular structure of the mitochondrial cavity, revealing axial mitochondrial widths at 118-150 nm. The half-widths at maximum of the axial localizations are at 21-39 nm (Figure 4b-4c). The high axial localization precision confirms the blinking kinetics of NOSR is exceedingly engineered through the photo-trigger self-blinking strategy, fulfilling the harsh requirements of three-dimensional super-resolution imaging.

**Figure 4.**
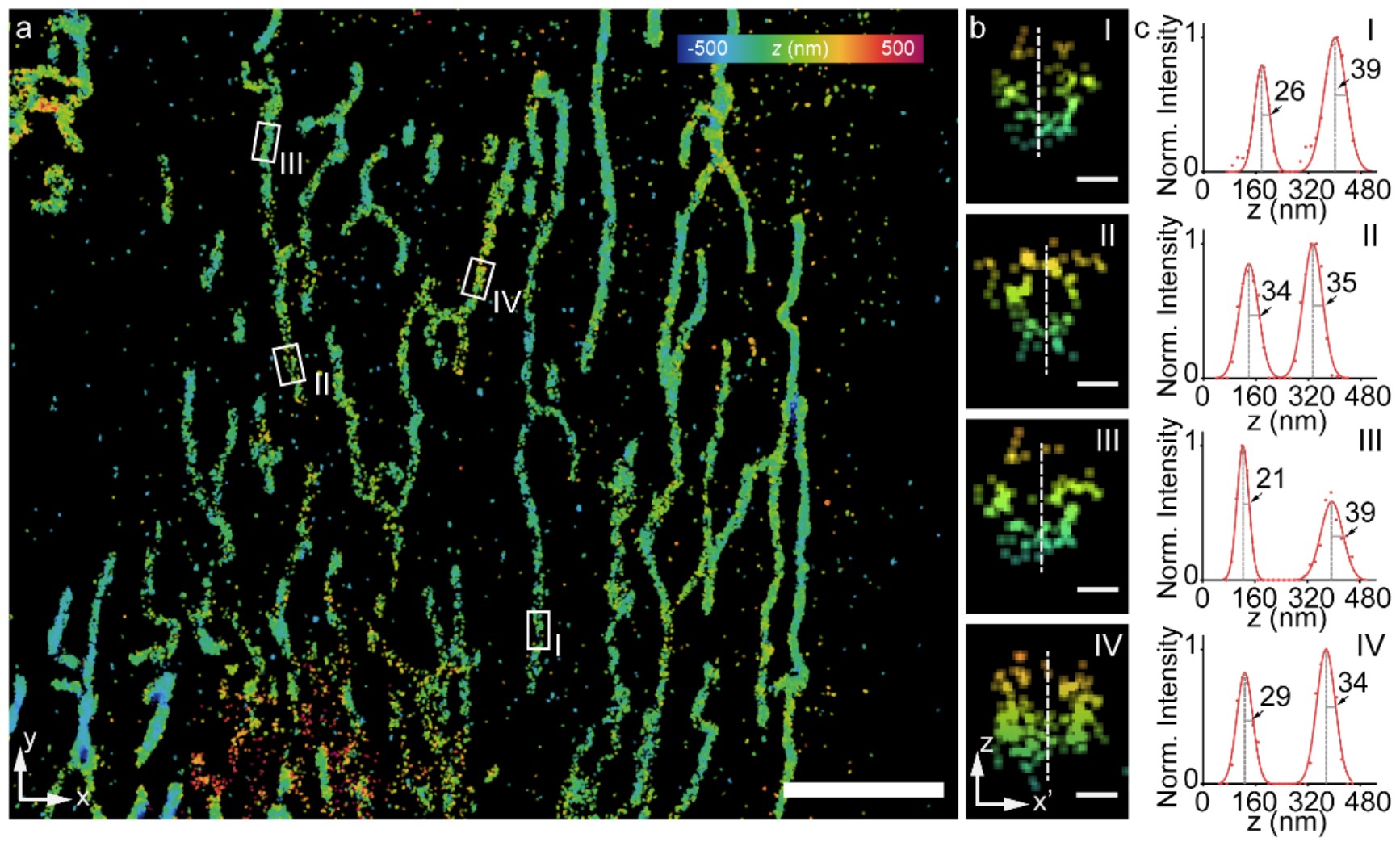
Three-dimensional single-molecule super-resolution imaging of mitochondrial outer membrane (a) with NOSR. (b) The x–z projection view of the white box in (a). (c) Intensity profile along the white dashed line in (b). Scale bars: 5 μm (a). 0.2 μm (b).

In conclusion, we explored the engineering of blinking kinetics through molecular structural design. A photo-trigger self-blinking fluorophore NOSR was built upon introduction of the nitroso-caging group into the self-blinking sulfonamide rhodamines matrix. Two modifications were strategically united without interference, as the photo-uncaging process released a self-blinking rhodamine. Single-molecule study revealed that the NOSR demonstrated a large number of self-blinking dark-bright events in a photo-gated fashion, presenting supremacy in the sparsity and durability over self-blinking analogues. In high-density labeled immunolabeling specimens, NOSR enabled robust imaging integrity for both two- and three-dimensional super-resolution imaging without any excess imaging enhancing buffer. We envision our photo-trigger self-blinking strategical molecular design pave a new road for developing dyes with controllable single-molecule blinking flexibility to satisfy the high demands from future super-resolution imaging.

## Acknowledgements

This work was supported by the Foundation of China (Nos. 22004011, 22174009 and 22374013). Fluorescent imaging was performed with the support of the Chemical Analysis and Research Center at Dalian University of Technology.

