## Supplement for "Photo-Uncaging Triggers on Self-Blinking to Control Single-Molecule Fluorescence Kinetics for Super-Resolution Imaging"

### 1 Experimental method

1.1 General

All regents were purchased from commercial suppliers and used as received. Column chromatography was performed with silica gel (200-300 mesh). $^1$H NMR and $^{13}$C NMR were measured on Bruker Avance II 400, Bruker AVANCE III 500 and Varian MERCURY 400 spectrometers. Mass spectra and high-resolution mass spectra were recorded on UPLC/Q-TOF Mass spectrometers.

1.2 Synthesis of NOSR

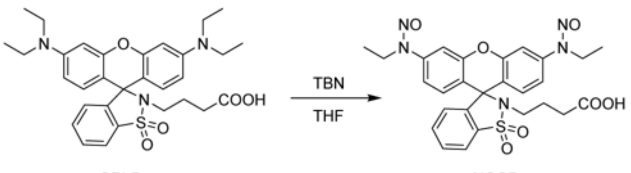

SRhB[1] (50 mg, 91 μmol) was dissolved in dry tetrahydrofuran (6 mL) and butyl nitrite (110 uL, 0.91 mmol) was added. The mixture was heated to 60 °C and stirred for 1 h. After the completion of the reaction, solvent was removed under reduced pressure. The crude product was further purified through silica gel column chromatography with a mixture of dichloromethane, ethyl acetate and acetic acid (100/2/1, v/v/v) as eluent. The collected solution was washed to remove acetic acid and then dried in vacuo to afford NOSR (35 mg, yield 68%). HRMS (ESI) *m*/*z* called for $C_{27}H_{28}N_5O_7S$ $[M+H]^+$: 566.1704; found: 566.1701 ($z$ = 1). $^1$H NMR (400 MHz, $CDCl_3$) δ 7.90 (d, J = 7.7 Hz, 1H), 7.52 (dt, J = 19.1, 7.4 Hz, 2H), 7.41 (d, J = 1.9 Hz, 2H), 7.25 (dd, J = 8.7, 2.0 Hz, 2H), 7.19 (s, 2H), 6.95 (d, J = 7.5 Hz, 1H), 3.98 (q, J = 7.2 Hz, 4H), 3.01 (t, J = 7.0 Hz, 2H), 2.17 (t, J = 7.0 Hz, 2H), 1.88 – 1.55 (m, 2H), 1.11 (t, J = 7.2 Hz, 6H). $^{13}$C NMR (101 MHz, $CDCl_3$) δ 177.15, 151.59, 143.87, 142.97, 134.12, 133.08, 130.70, 130.08, 126.22, 120.92, 118.01, 114.57, 106.55, 65.03, 40.18, 38.55, 30.10, 23.28, 11.73.

#### 1.3 Comparison fluorophores

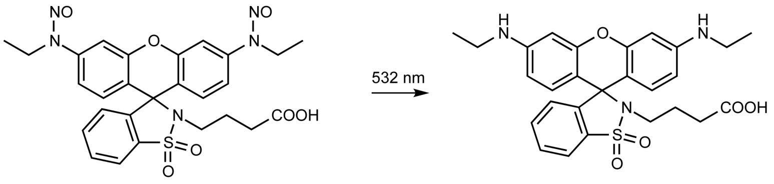

NOSR SRhod

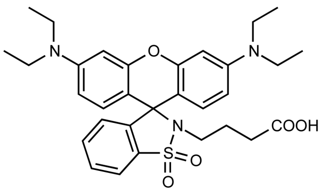

SRhB

#### 1.4 Spectral measurement

Absorption spectra was recorded with an Agilent UV-Vis absorption 8453 spectroscopy system and the fluorescent spectra was recorded with an Agilent Cary Eclipse fluorescence spectrophotometer.

For solvents utilized in spectra analysis, PBS is phosphate-buffered saline (10 mM, pH 7.3), prepared from MilliQ water (filtered with 0.22 µm syringe filter units). Other solvents were spectrophotometric grade and utilized as received.

*The photoactivation test and photoconversion quantum yields test.* The photoconversion quantum yields were determined following the method described in a reference.[2] The test solvents were PBS (contained 10% MeOH). The sample was saturated with argon and measured in a 4 mL quartz cuvette. A portable 365 nm lamp (~540 µW/cm$^2$, measured by Lianhuicheng LH-127 light power density meter) was utilized for photoactivation.

*Relative fluorescence quantum yield.* To minimize the effect of reabsorption/reemission, all testing samples were diluted with a visible absorption peak less than 0.09. MPR (previous determined by absolute quantum yield spectrometer) in water was deployed as the reference standard.[3] Identical excitation wavelength was utilized for both standard and samples. Their fluorescence was recorded under identical configurations. The relative quantum yield (precision: ± 10%) of a sample was calculated according to the equations below:

$$\Phi_s = \Phi_{st} \times \frac{A_{st}}{A_s} \times \frac{I_s}{I_{st}} \times \frac{n_s^2}{n_{st}^2} \quad (1)$$

Φ was the quantum yield. A was the absorption at the excitation wavelength. I was the integrals of fluorescence. n was the refractive index of the solvents. The subscripts of these elements were listed as: $_s$ was the sample and $_{st}$ was the standard (reference sample).

*The pK$_{cycling}$ test.* The photoactivation product of NOSR were prepared as 2 µM solution for measurements. The buffer was PBS containing 10% ethanol. The proton responding curve was obtained through pH titration of the fluorophore solution.

pK$_{cycling}$ was calculated through Henderson–Hasselbalch equation,

$$I = \frac{1}{b(10^{pH-pKa+1})} + c \quad (2)$$

*I* was the integration of fluorescence peak or absorption peak. b and c were constants.

#### 1.5 Microscopy

Single-molecule and Super-resolution imaging were studied with a total internal reflection fluorescence microscope (TIRFM) built on an Olympus IX71 inverted microscope as described earlier.[3,4] The applied laser was a 200 mW 532 nm continuous laser (Coherent, Sapphire 532-200). The irradiation power was 2.0 kW/cm$^2$ for imaging. The laser angle was adjusted to highly inclined illumination mode. The deployed objective was a 100× TIRF oil objective (UAPON, Olympus; 1.49 numerical aperture). The signals were recorded on an EMCCD (iXon DU-897U, Andor). For 3D super-resolution imaging, a cylinder lens (f = 1557.42 mm) was inserted into the fluorescence emission path to induce astigmatism for 3D super-resolution imaging.

#### 1.6 Cell culture

HeLa (helacyton gartleri) cells were purchased from Cell Bank of Type Culture Collection of Chinese Academy of Sciences. The cells were cultured in full growth medium (cell culture media), i.e. minimum Eagle's medium (MEM) supplemented with 10% fetal bovine serum (FBS, Hyclone). The cells were cultured in humidified atmosphere at 37 °C and 5 % $CO_2$. Cells were seeded on 25 mm cover slip (Fisherbrand, 12-545-102) for 1–2 days to reach 70-90% confluence before imaging.

#### 1.7 Antibody labeling

Fluorophores were labeled to proteins through two steps: (1) activation through an NHS (N-hydroxysuccinimide) ester; (2) covalent coupling to the α-amine groups of proteins. The labeling method following a procedure described in our previous report.[2,3]

#### 1.8 Fixation and labeling of microtubules and mitochondrial outer membrane in fixed cells

*Fixation.* In the preparation of microtubules, HeLa cells were first pre-extracted with BRB buffer (0.25% Triton X-100, 80 mM PIPES, pH = 6.8, 1 mM $MgCl_2$ and 1 mM EGTA) supplemented with 4 mM EGTA for 10 s. Then these cells were fixed with 0.5% glutaraldehyde in BRB buffer at room temperature for 10 min. In the preparation of mitochondrial outer membrane, HeLa cells were first fixed with 37 °C pre-warmed 3% PFA and 0.5% GA in PBS at room temperature for 15 min. The background fluorescence from fixation was quenched with fresh 0.1% $NaBH_4$ (PBS solution) for 7 min. Before immunostaining, the remaining fixation solution was washed out through PBS rinse for three times.

*Immunostaining.* Labeling microtubules, HeLa cells were stained with a rabbit polyclonal antibody against α-tubulin (Beyotime) at 1:100 dilution in 30% NGS (normal goat serum, Beyotime, C0265) for 12 h at 4 °C. Labeling mitochondrial outer membrane, HeLa cells were washed three times with PBS and then treated with blocking buffer (3% BSA and 0.2% Triton X-100 in PBS) for 1 h, gently rocked at room temperature. Then cells were stained with a rabbit polyclonal antibody against TOMM20 (Beyotime) at 1:100 dilution in 30% NGS (normal goat serum, Beyotime, C0265) for 12 h at 4 °C. The unbounded antibodies were washed out through PBST (PBS contained 0.1% Tween-20) rinse for three times. Then these cells were stained with fluorophore labeled secondary antibodies at 4 °C for 1 h. The unbounded antibodies were washed out through PBST rinse for three times.

#### 1.9 Single-molecule imaging

Antibodies labeled with fluorophores were freshly diluted in PBS at a low concentration to minimize the overlapping between different molecules and transferred to the surface of clean coverslips. The adhesion of antibodies to the surface proceeded for 30 s and the unbounded proteins were washed out through three times rinses of PBS. SRhod was obtained through uncaging of NOSR through 2 s irradiation of ~100 W/cm$^2$ 375 nm laser light. The integration time of single-molecule signals was 10 ms and 6000 frames were totally recorded for each measurement. At least 5 measurements were performed for each irradiation condition of every fluorophore.

The integration time of the single molecule signal was 10 ms, and a total of 10000 frames were recorded for measuring the temporary shift of NOSR.

1.10 Post-processing of super-resolution imaging data.

Single-molecule data were automatically processed with a home-written Matlab software as described in our previous report.[1] Briefly, a wavelet filter was utilized to remove background noises on raw image stacks. Single-molecule candidates were identified from filtered image stacks and their single-molecule fluorescent trajectories were further extracted from unfiltered raw data. These trajectories were analyzed with a hidden Markov model to obtain the state transition trajectories through Baum–Welch, forward-backward and Viterbi algorithms (A gaussian distribution model was deployed for the observing probability distribution). Single-molecule photophysics were measured from single-molecule trajectories and their state transition trajectories.

*Brightness.* Single-molecule brightness in this manuscript was determined as the photon counts from a single molecule during the acquisition time of single frame (10 ms). In other words, the photons emitted from a molecule during 10 ms acquisition time.

*On time.* The time duration of a bright state.

*Dark time.* The time duration of a dark state.

*Switch numbers.* The counted times of bright-to-dark transitions of a molecule.

*Duty cycle.* Duty cycle was defined as the ratio between the total time of a single molecule spent on its on-state and the entire imaging time:

$$\text{Duty Cycle} = \frac{\sum t_{on}}{T} \quad (3)$$

$t_{on}$ was the dwell time of a switching event. $T$ was the entire imaging time (100 s in this study).

1.11 Super-resolution imaging

Ambient light should be avoided during the entire staining procedure, to minimize the accidental photo-uncaging of NOSR. Before imaging, the media was replaced to PBS for fixed cells. A conventional image was acquired with low laser intensity before the super-resolution imaging. During super-resolution imaging, a continual 532 nm laser (2 kW/cm$^2$) was utilized for excitation. The single-molecule photoswitching signals were recorded at 25 Hz.

1.12 Post-processing method

Super-resolution imaging analysis was performed in home-written Matlab codes.[5,6]

#### 2 Supplemental figures

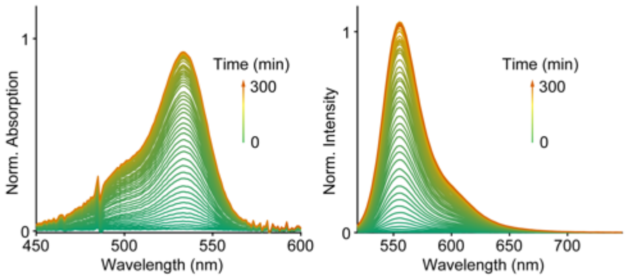

Figure S1. Normalized absorption and emission spectral changes of NOSR during photoactivation under 365 nm light irradiation in PBS (contained 10% MeOH).

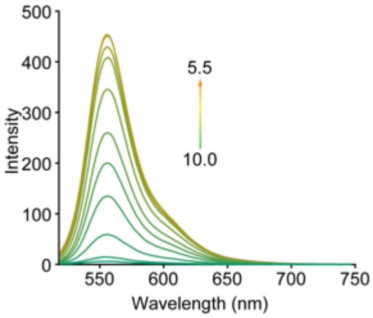

Figure S2. Fluorescence spectra of the photoactivation product of NOSR at a series of pH conditions.

Table S1. Single-molecule Photophysics of NOSR, SRhod and SRhB

| Dye | Laser power ($W/cm^2$) | Brightness (photons/10ms) | Duty cycle | Switch numbers | On time (ms) | Dark time (s) | Bleach time (s) |
|---|---|---|---|---|---|---|---|
| NOSR | 500 | 92 ± 3 | 0.017±0.004 | 21.6±3.8 | 45±3.0 | 0.8±0.16 | 17.4±1.2 |
| | 1000 | 154±13 | 0.011±0.003 | 18.4±3.7 | 36±2.3 | 1.0±0.06 | 17.6±3.1 |
| | 1500 | 227±17 | 0.008±0.001 | 16.8±0.7 | 29±2.4 | 1.1±0.05 | 18.1±0.5 |
| | 2000 | 351±52 | 0.006±0.0004 | 15.1±1.0 | 24±0.1 | 1.2±0.06 | 17.3±0.6 |
| SRhod | 500 | 114±4 | 0.028±0.003 | 29.7±2.9 | 57±4.8 | 0.7±0.07 | 22.1±2.8 |
| | 1000 | 172±17 | 0.015±0.002 | 23.5±2.2 | 39±2.8 | 0.9±0.07 | 20.6±1.5 |
| | 1500 | 217±20 | 0.010±0.001 | 19.0±1.5 | 32±2.3 | 1.1±0.06 | 19.6±0.6 |
| | 2000 | 353±14 | 0.007±0.001 | 15.4±1.0 | 27±2.4 | 1.3±0.02 | 15.6±1.5 |
| SRhB | 500 | 89±6 | 0.047±0.010 | 25.9±2.4 | 107±24 | 0.6±0.13 | 18.4±3.0 |
| | 1000 | 123±6 | 0.023±0.007 | 14.4±1.1 | 96±29 | 0.8±0.15 | 12.2±1.8 |
| | 1500 | 195±12 | 0.015±0.002 | 11.4±2.9 | 79±20 | 0.7±0.14 | 8.1±1.4 |
| | 2000 | 231±24 | 0.011±0.001 | 9.7±1.4 | 67±15 | 0.8±0.18 | 7.5±1.5 |

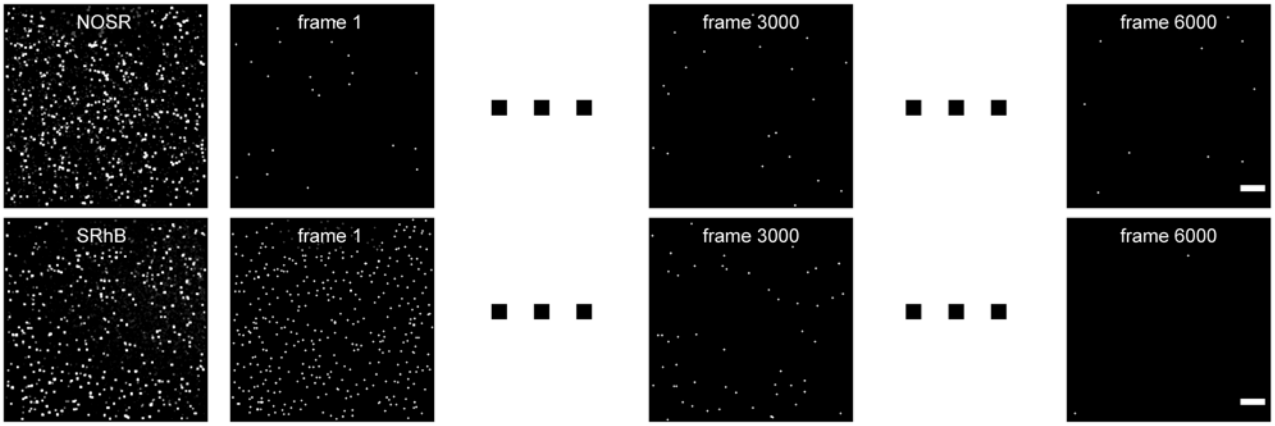

Figure S3. The molecular distribution density of NOSR and SRhB in single-molecule imaging with high density labeling changes with time. Scale bars: 5 μm.

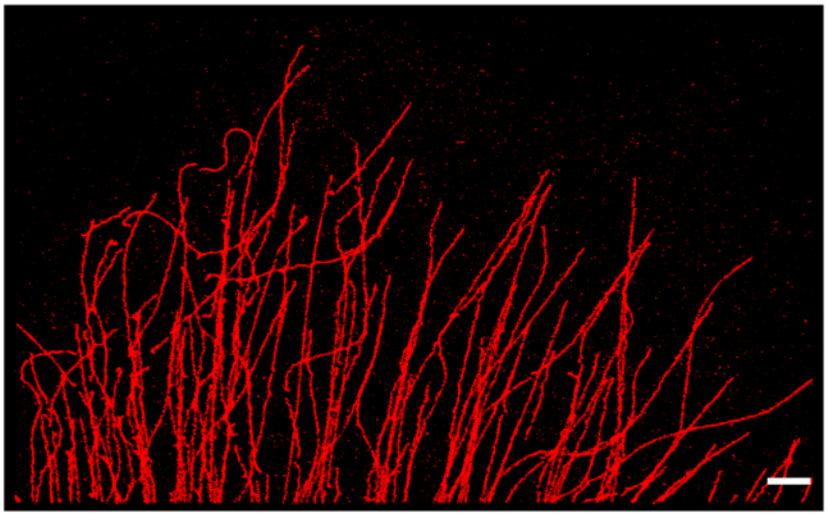

Figure S4. Super-resolution imaging of microtubules in a fixed HeLa cell with primary antibody against α-tubulin and secondary antibody conjugated to NOSR. Scale bar: 2 μm.

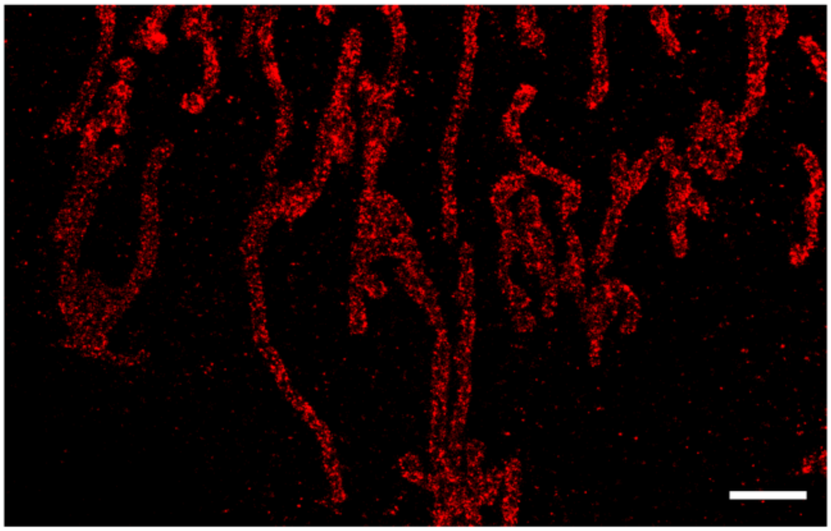

Figure S5. Super-resolution imaging of mitochondrial outer membrane in a fixed HeLa cell with primary antibody against TOMM20 and secondary antibody conjugated to NOSR. Scale bar: 2 μm.

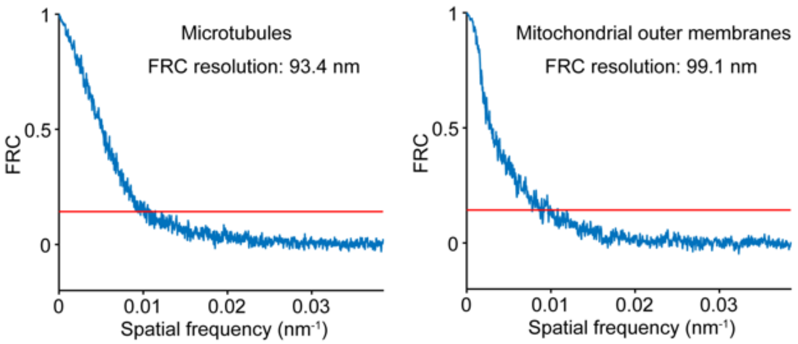

Figure S6. Fourier ring correlation curve analysis of reconstructions from NOSR in Figure 3.

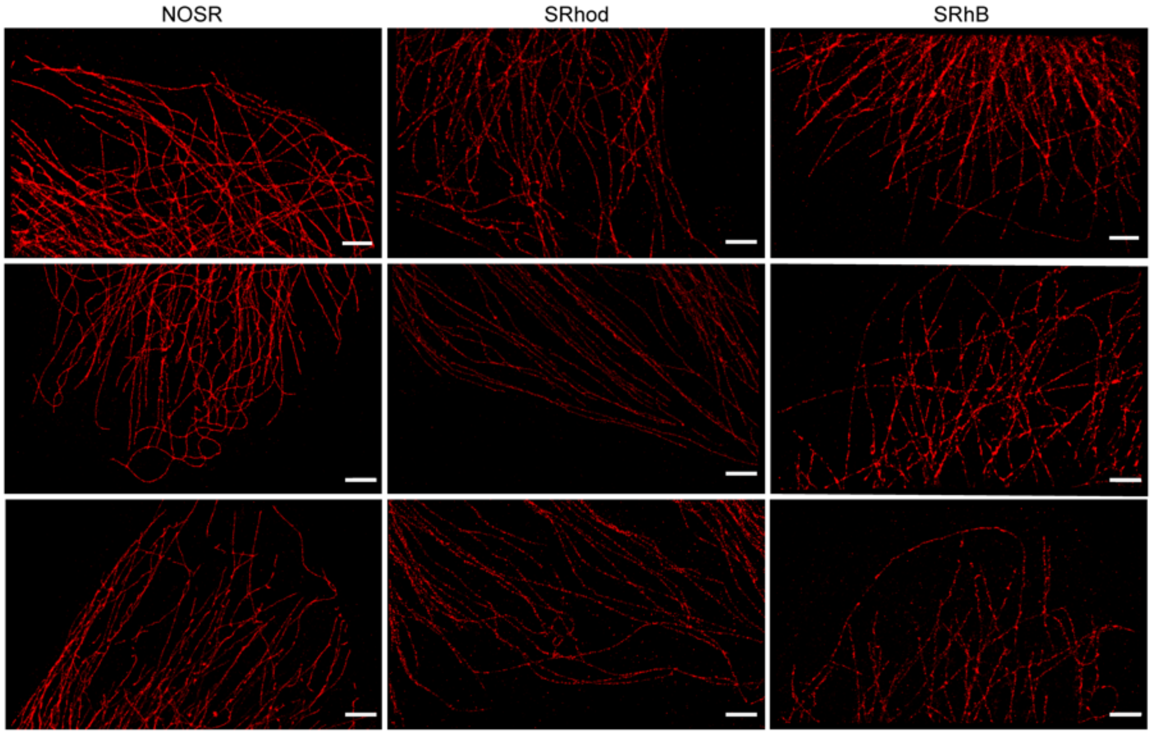

Figure S7. Comparison on the super-resolution imaging of microtubules in fixed cells results between NOSR, SRhod and SRhB. Scale bars: 2 μm.

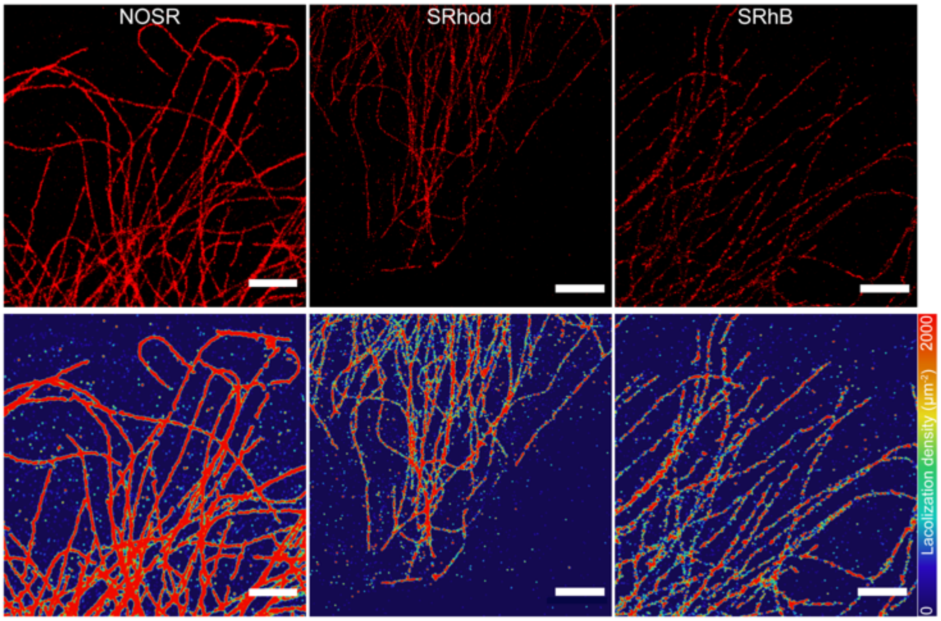

Figure S8. Comparison on the super-resolution imaging results and the corresponding localization density maps of microtubules between NOSR、SRhod and SRhB. Scale bars: 2 μm.

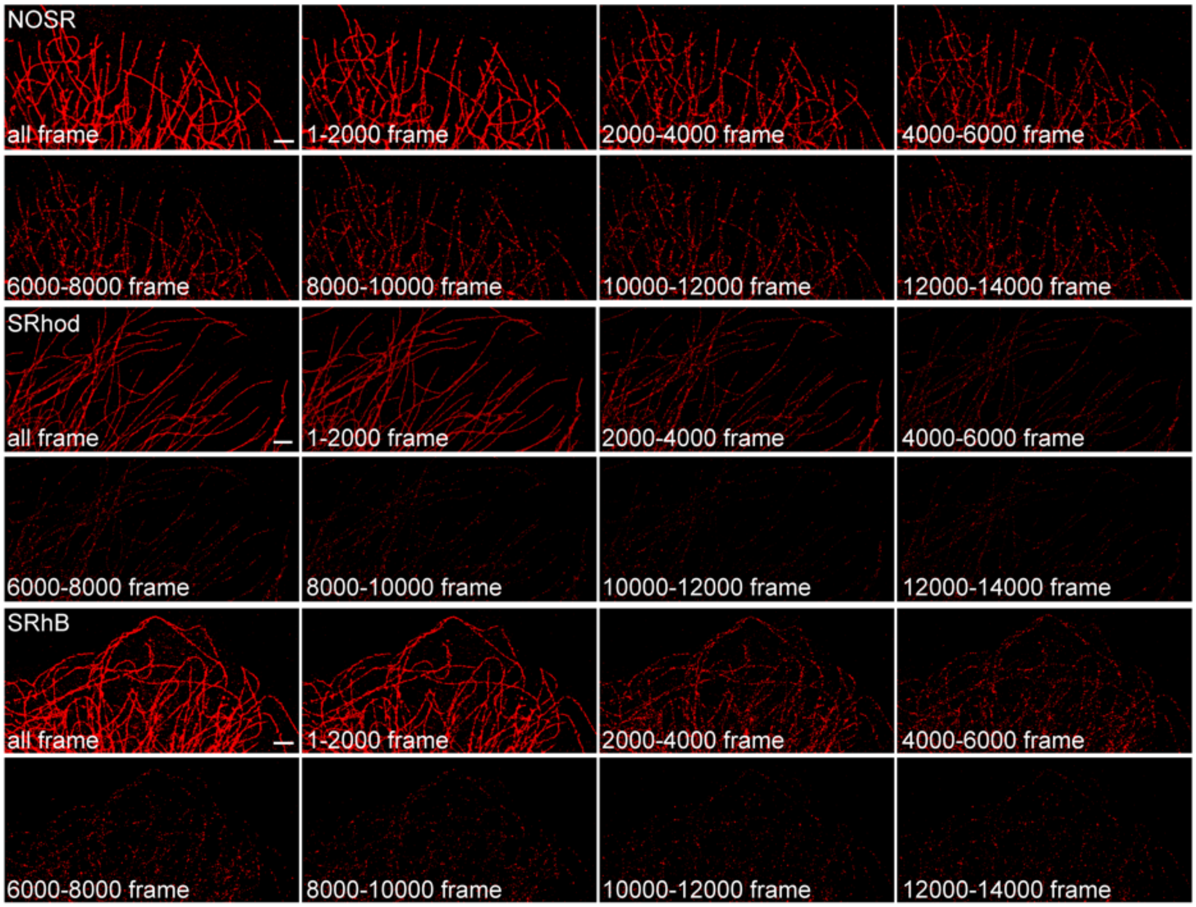

Figure S9. Comparison on the super-resolution imaging of microtubules during different imaging times between NOSR, SRhod and SRhB. Scale bars: 2 μm.

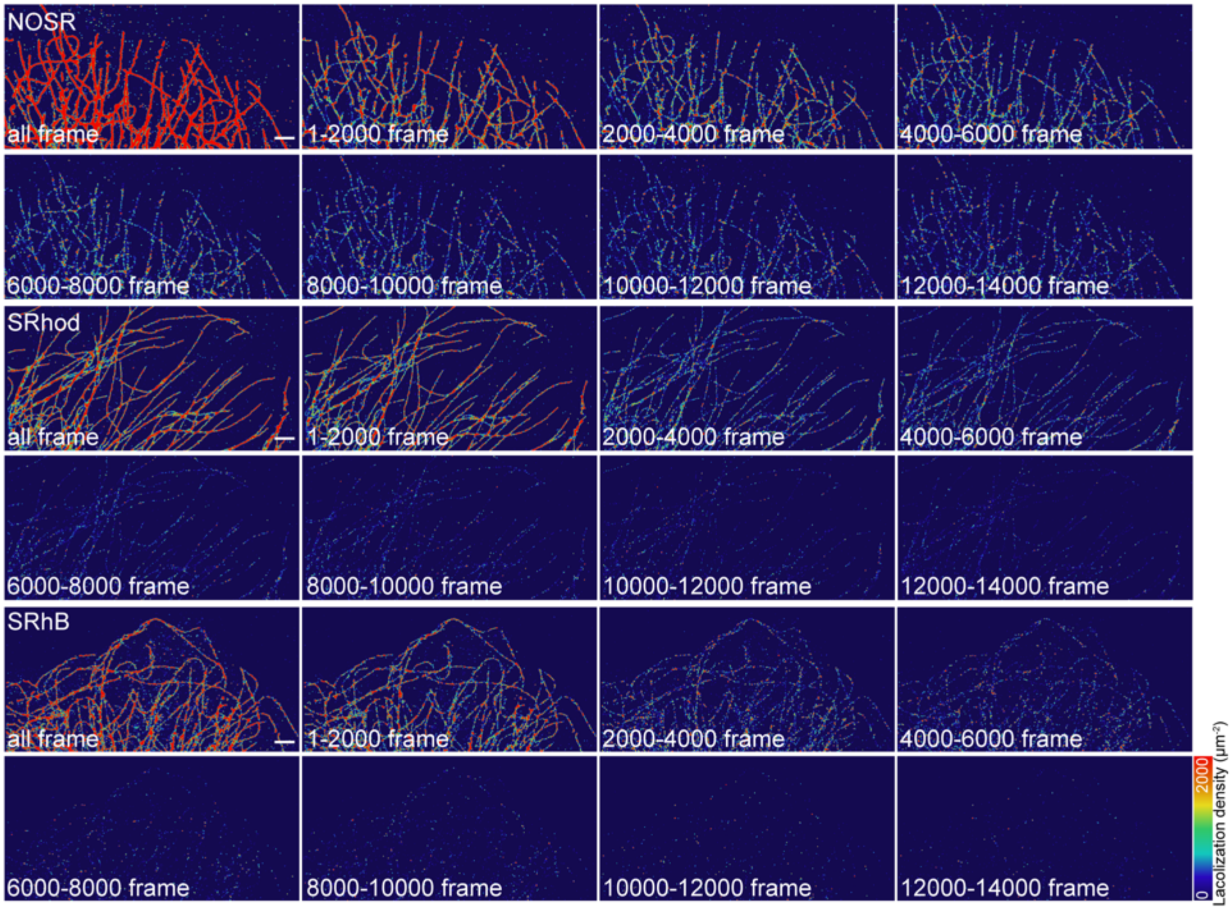

Figure S10. Comparison on the corresponding localization density maps of microtubules in Figure S9. Scale bars: 2 μm.

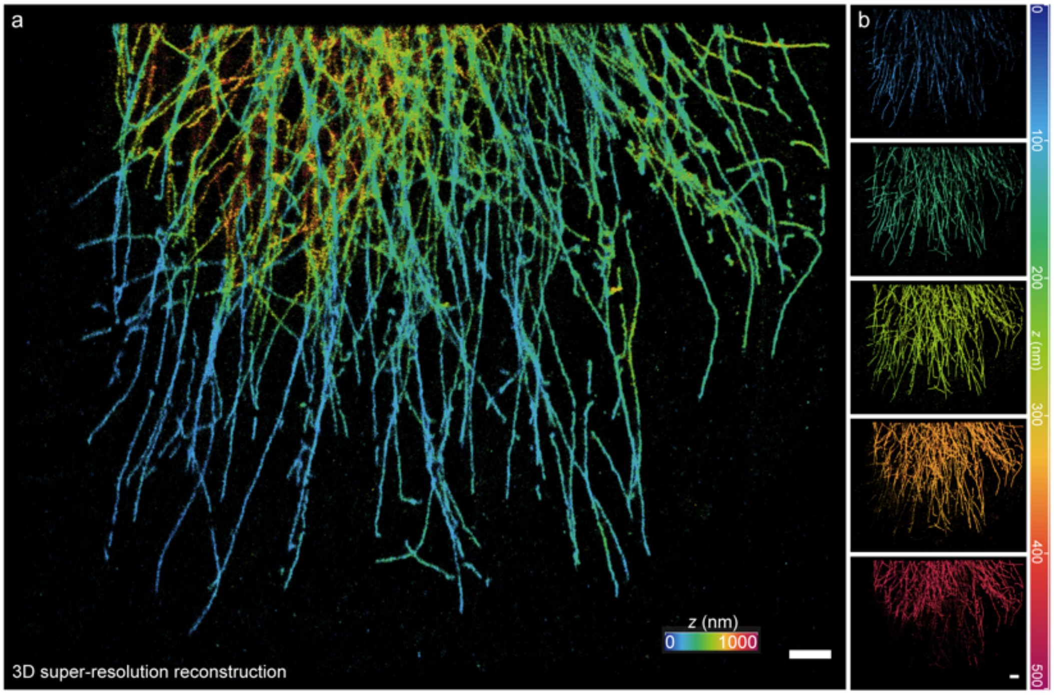

Figure S11. (a) Three-dimensional SMLM imaging of microtubules in a fixed cell by NOSR. (b) Serial Z-step 3D reconstructions. Scale bars: 2 μm.

### 3 Characterization spectra

3.1 $^{1}$H NMR spectrum of NOSR

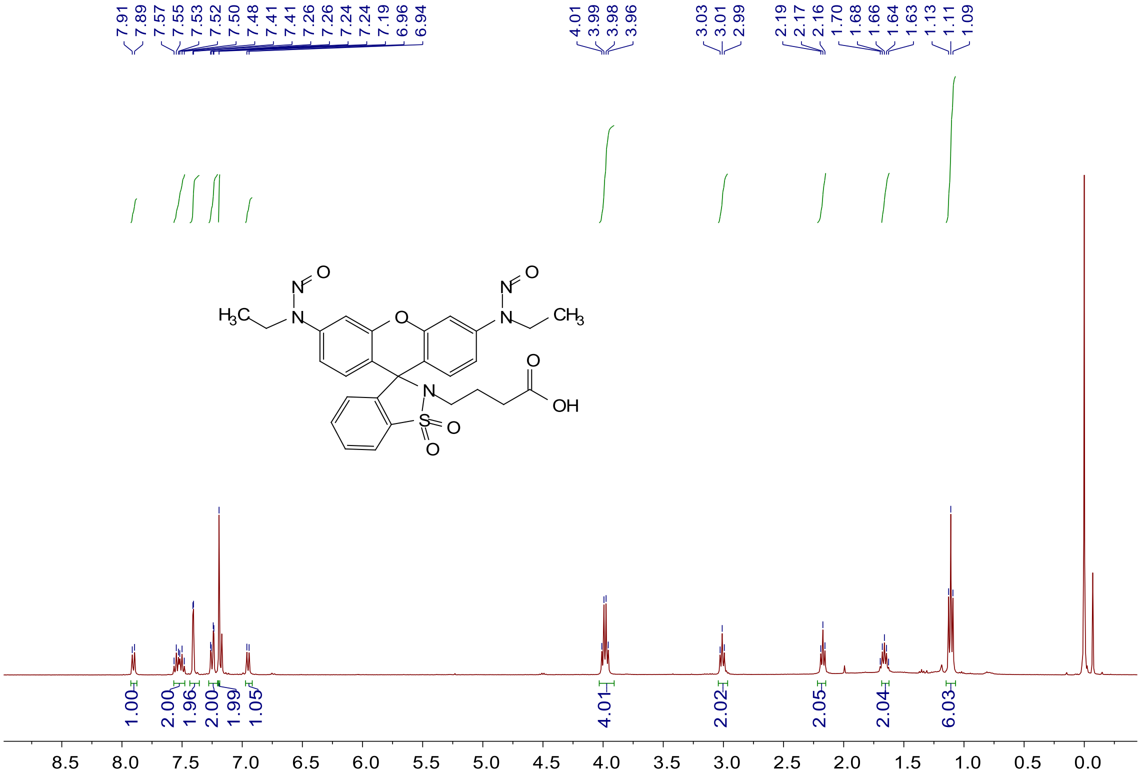

3.2 $^{13}$C NMR spectrum of NOSR

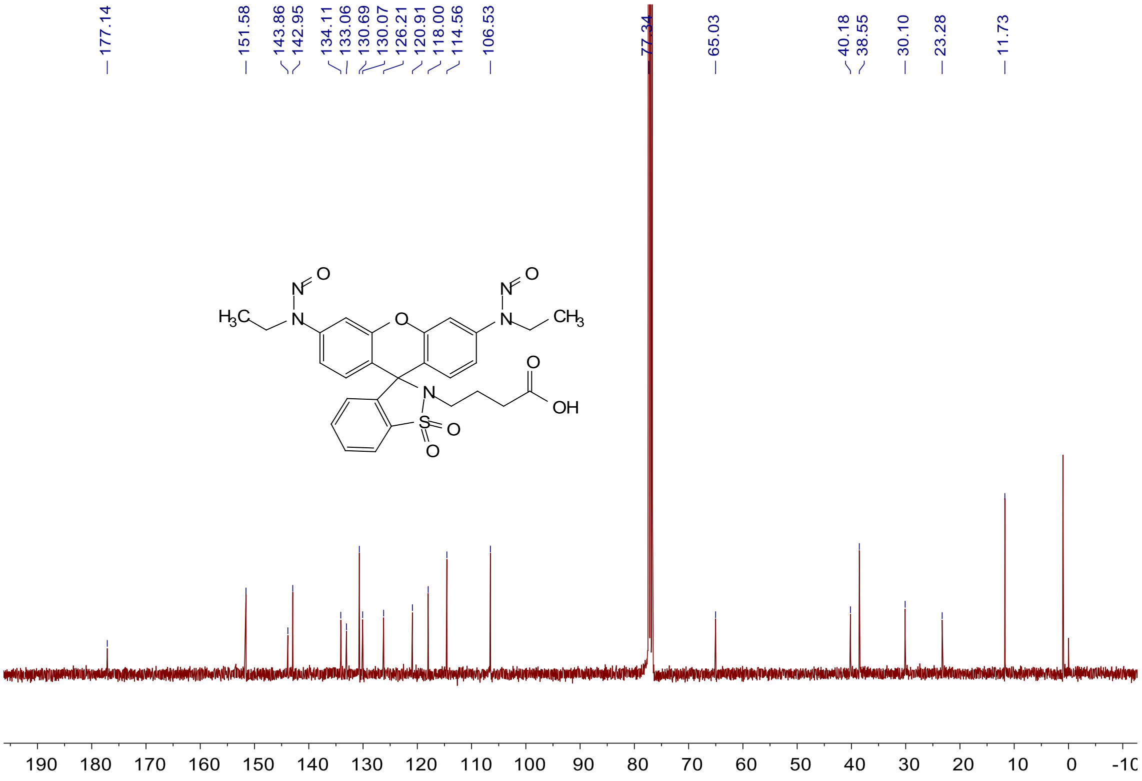
